# Stronger density-dependent growth of Japanese sardine with lower food availability: Comparison of growth and zooplankton biomass between a historical and current stock-increase period in the western North Pacific

**DOI:** 10.1101/2021.12.26.474216

**Authors:** Yasuhiro Kamimura, Kazuaki Tadokoro, Sho Furuichi, Ryuji Yukami

**Author notes:** **Corresponding author:** Y. Kamimura.

## Abstract

Density dependence is a fundamental concept for fish population dynamics. Although density-dependent growth and maturity among older juveniles and adults is important for regulating fish population size and for fisheries management, the mechanism of density dependence for marine fishes remains unclear. Here, we examined changes in Japanese sardine growth with increasing abundance beginning in the 2010s and how the current pattern of density-dependent growth differs from that of a previous stock-increase period from the 1970s to early 1980s. During the current period of increasing abundance, mean standard length has already dropped to the lowest level yet observed and growth has declined more sharply with increased abundance than in the 1970s and 1980s. Mesozooplankton biomass in July in the summer feeding grounds was also lower during the current period. Therefore, our results suggest that summer food availability in the western North Pacific controls the strength of density-dependent growth. A lower carrying capacity for Japanese sardine could account for the stronger density dependence of growth observed in the 2010s; this indicates that future Japanese sardine abundance might not increase as much as in the 1980s unless food availability improves.

## 1. Introduction

Density dependence plays a key role in population dynamics. Changes in mortality, growth, maturity, and habitat selection during population fluctuations have been observed in many fishes (e.g., Lorenzen and Enberg, 2002; Rose et al., 2001; Zimmermann et al., 2018). When constructing population dynamics models for fisheries management, it is common to consider density dependence in early life stages (e.g., for egg production or for the period between hatching to recruitment) as a stock– recruitment relationship (Beverton and Holt, 1957; Ricker, 1954). On the other hand, density dependence in the late juvenile and adult stages (e.g., on growth and maturation) is rarely considered in these models because the process is currently not fully understood (Andersen et al., 2017). Density-dependent growth and maturity in later life stages could have important implications for fish population regulation (Lorenzen and Enberg, 2002) and for calculating reference points for management (Van Gemert and Andersen, 2018). Therefore, there is an urgent need to clarify the underlying processes to improve the implementation of effective management schemes.

For lake- and stream-dwelling fishes such as salmonids, the theoretical framework for the density dependence of growth and mortality (i.e., decreases in growth or increases in mortality with increasing population density) has been examined in detail through field and laboratory experiments as well as field observations (e.g., Matte et al., 2020a, 2020b). These studies have demonstrated the importance of food availability in the process of density dependence (Amundsen et al., 2007; Jenkins et al., 1999; Post et al., 1999), and of the application of both density-dependent and independent factors to growth models (Matthias et al., 2018). However, for marine fishes, the frameworks and processes of density dependence have not yet been fully examined (Andersen et al., 2017), despite the many studies that have reported negative density dependence in growth (e.g., Rueda et al., 2015; Zimmermann et al., 2018). Examining how the density-dependent growth of marine fishes is altered by environmental conditions (i.e., food availability) could prove useful for understanding the processes underlying population fluctuations and effective management schemes.

The Japanese sardine *Sardinops melanostictus* Temminck & Schlegel, 1844, supports one of the largest fisheries of small pelagic fishes in the western North Pacific, and its abundance has fluctuated repeatedly on a multidecadal scale (Furuichi et al., 2020a; Kuwae et al., 2017). These fluctuations are particularly evident in the history of the Pacific stock of Japanese sardine over the past 50 years. The stock initially increased in the early 1970s (Wada and Jacobson, 1998) and maintained high recruitment into the 1980s (Fig. 1), with total biomass over 13 × 10^6^ t (Furuichi et al., 2020a), until the stock collapsed due to recruitment failure after 1988 (Watanabe et al., 1995). Abundance remained at low levels until 2010, after which the stock has steadily recovered. This recent trend toward increasing abundance is likely to be similar to that in the 1970s based on estimates reported in a stock assessment (Furuichi et al., 2020a; Fig. 1) and in a previous study (Wada and Jacobson, 1998).

**Fig. 1.**
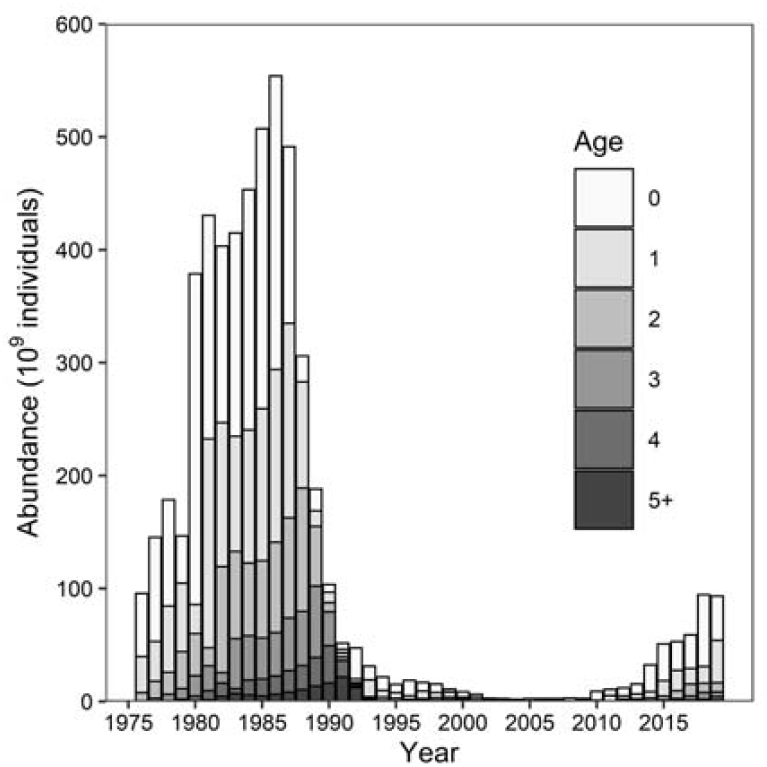
Abundances (10^9^ individuals) of the Pacific stock of Japanese sardine estimated by Furuichi et al. (2020a).

Growth patterns of Japanese sardine are clearly affected by intra-specific density dependence (Kawabata et al., 2011; Kim et al., 2006; Morimoto, 2003; Ohshimo et al., 2009). Growth and maturity-at-age of the Pacific stock decreased in the late 1970s to early 1980s, when abundance was increasing, but increased in the 1990s, when abundance was in decline (Kawabata et al., 2011; Morimoto, 2003). For the most recent period of increasing abundance (i.e., after 2010), although recent studies have reported annual changes in larval and juvenile growth (Furuichi et al., 2020b; Niino et al., 2021), late-juvenile and adult growth patterns remain unexamined. This information is urgently needed to inform management strategies and forecasts of future abundance for this stock.

As mentioned above, although the commercial importance and growth variability of Japanese sardine have interested many researchers, the annual periodicity of otolith increments has not been confirmed in this species, meaning that scales are the only available method for aging individuals (e.g., Nakai, 1962). This method has been used successfully as a basis for stock assessment (Furuichi et al., 2020a) as well as in several studies of late-juvenile and adult growth (Kawabata et al., 2011; Kim et al., 2006; Morimoto, 2003; Wada and Jacobson, 1998). However, scale-based aging can be somewhat unreliable as scales can easily detach from the body and be replaced by regenerated scales. Generally, mean otolith ages tend to be slightly more precise (Campana, 2001), and otolith-based age determination is the dominant method for other sardines (Fletcher and Blight, 1996; ICES, 2019; Yaremko, 1996). Therefore, there is an urgent need to confirm the annual periodicity of otolith increment formation in Japanese sardine.

In the present study, we aim to understand the processes underlying the density dependence of growth in the Pacific stock of late-juvenile and adult Japanese sardine, with a particular focus on food availability. To do so, we relied partly on datasets of changes in Japanese sardine growth from 1976 to the early 1980s (Wada and Kashiwai, 1991a, 1991b) and of zooplankton biomass since the 1970s in Oyashio waters in summer (Odate, 1994; Tadokoro et al., 2005), which is when Japanese sardine undergo northward feeding migrations (Nakai, 1962; Yatsu, 2019). First, we investigated changes in growth during the period of increasing abundance since 2011 by confirming the annual periodicity of otolith-ring formation. Then, we compared Japanese sardine growth and summer zooplankton biomass during the period of stock increase in the 2010s (hereafter, “the current period of increase”) against the period of increase from the 1970s to early 1980s (hereafter, “the previous period of increase”) with the assumption that summer food availability affects the strength of density dependence. Finally, we briefly discuss the implications of our results for the future trajectory of the Pacific stock of Japanese sardine.

## 2. Material and methods

### 2.1. Fish sampling

Japanese sardine samples were selected from commercial catches landed at Hachinohe port (40°5′N, 141°6′E) by purse seine fisheries and from among individuals collected during scientific surveys. All samples were caught from 36–48°N and 141–172°E in the western North Pacific from 2011 to 2020 (Fig. 2). Scientific surveys were conducted by using a surface–midwater trawl net (NST-99, Nichimo Co., Ltd.; opening dimensions: 30 m × 30 m; cod-end mesh size: 17.5 mm) towed once at each station (15–60 min at 3.0– 5.0 kn at 0– 30-m depth) from aboard the training vessel (T/V) *Hokuho Maru*. Samples were frozen at −20 °C on board for subsequent measurement and otolith analysis. Standard length (SL) was measured to the nearest 0.1 mm by using digital calipers in the laboratory.

**Fig. 2.**
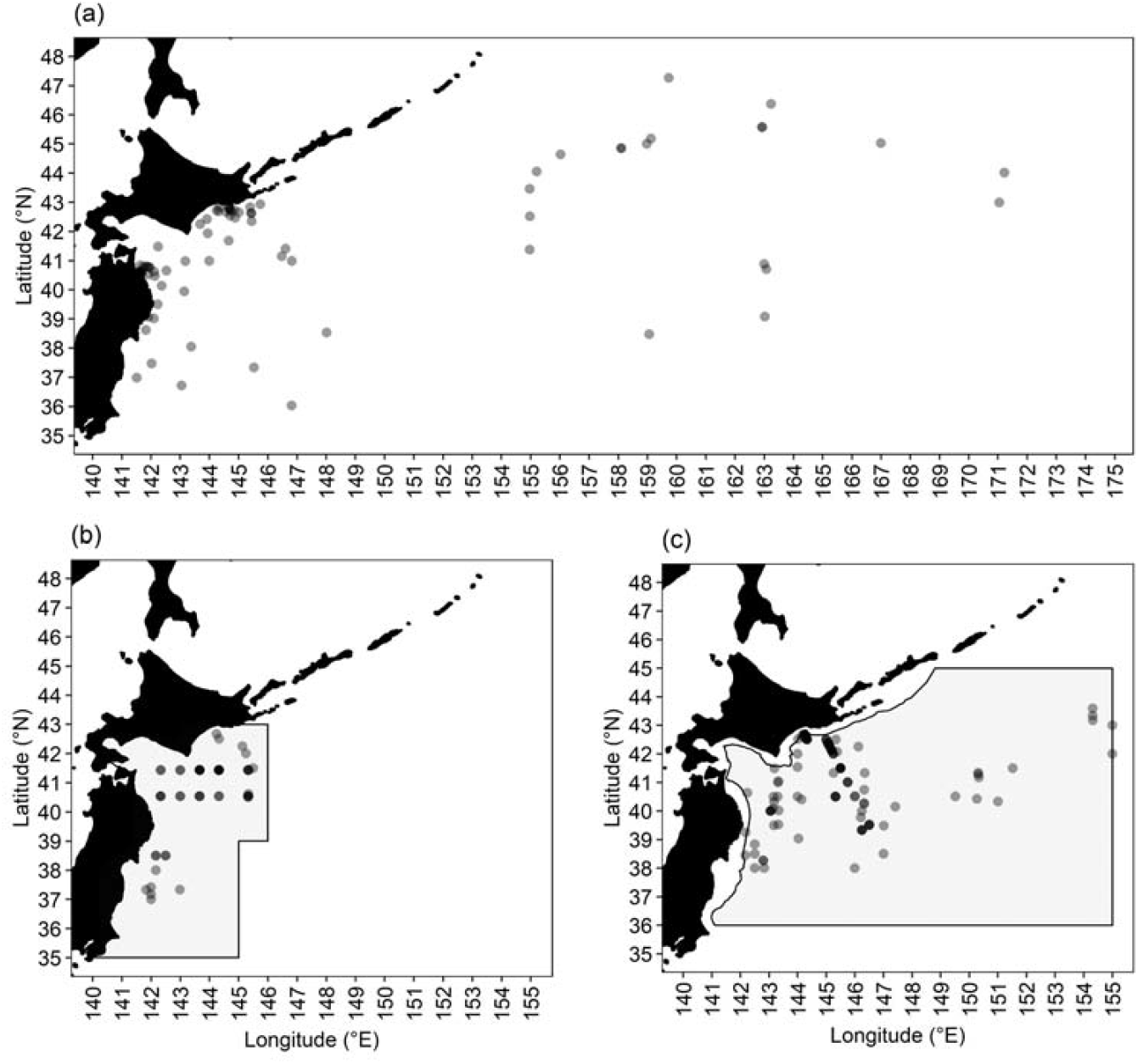
Sampling locations of (a) Japanese sardine, (b) zooplankton in June, and (c) zooplankton in July. Shading indicates the areas defined by Odate (1994) and Tadokoro et al. (2005).

### 2.2. Age determination from sagittal otoliths

We randomly selected 5–50 individuals from each sampling day or station. Sagittal otoliths were extracted from each fish and washed with fresh water. The external faces of sagittae were observed on a glass petri dish filled with 99% ethanol against a black background under a stereo microscope (SZX16, Olympus Corp., Tokyo, Japan) with reflected illumination units, and photographs were captured by using a digital camera (DP27, Olympus Corp., Tokyo, Japan). We evaluated our otolith-based age determination by using edge analysis, a commonly used validation method, in which we recorded whether an opaque or translucent zone appears on the outer margin of an otolith during every month or season (Campana, 2001). The annual periodicity of otolith increment formation was confirmed (see results), and therefore each sample was aged by counting the number of translucent zones on the posterior side of the otolith; the annulus for a single year consisted of one opaque zone and one translucent zone. The hatch date was assumed to be 1 January for the purposes of age determination in this study. A previous report recommended the use of a reference radius to aid the identification of the first annual ring for sardines (ICES, 2019). Therefore, we measured the radius from the nucleus to the first translucent zone on the posterior side of the otolith (*r*_*1*_) to the nearest 1 µm for several selected samples of each age.

### 2.3. Growth estimation

Changes in growth of Japanese sardine during the previous period of increase (i.e., for the 1976 to 1982 year-classes [YCs] except for the 1979 YC), have been previously estimated by using the von Bertalanffy growth formula (VBGF) (Wada and Kashiwai, 1991a, 1991b). Therefore, we fit a VBGF to data for the 2011 to 2018 YCs to compare growth rates between the previous and current periods of increase. The VBGF was fitted separately to length-at-age data for each year-class to estimate cohort-specific growth parameters:

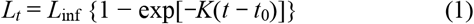

where *L*_*t*_, *L*_inf_, *K*, and *t*_0_ are SL at age *t*, asymptotic SL, growth coefficient, and hypothetical age at zero SL, respectively. Non-linear least-squares regression was used to estimate these parameters. Wada and Kashiwai (1991b) estimated VBGFs for samples collected from July to October and assumed a hatch month of September; hence, we modified the *t*_0_ of VBGFs obtained in their study by assuming a hatch month of January to better compare growth rates between the two periods.

The VBGF parameters for the previous period of increase were based on scale-based age determination. Because we used otolith-based age determination for the current period of increase, we evaluated the possible effects of the different choices of aging character by roughly comparing VBGFs estimated by otolith and scale data from the 2013 and 2018 YCs. Scale-based length-at-age was evaluated with a monthly age– length key (ALK) estimated from individuals caught on the Pacific side of northern Japan, which was the same as the ALK used for a previous stock assessment (Furuichi et al., 2020a). Since the ALK used 5-mm length intervals, the minimum SL was set to 2.5 mm. Scale-based VBGF parameters were estimated in the same way as for the otolith-based VBGF.

### 2.4 Examination of density dependence in growth

To examine the density-dependent growth of Japanese sardine, Kawabata et al. (2011) examined the relationship between body length at age estimated from a VBGF (*L*_*t*_) and *N*_*c*_, an index of the abundance that a given year-class experienced. In this study, we adopted the same index to examine density-dependent growth. Conceptually, *N*_*c*_ is the cumulative sum of Japanese sardine abundances over the period from when a given year-class is born to when it reaches the age *t* – 1, or in equation form: 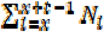, where *N*_*i*_ is the total population abundance in year *i*. The age range of samples previously used to estimate VBGF parameters during the previous period of increase was 1–4 (Wada and Kashiwai, 1991b); therefore, the range of *t* used in our study was 2–4.

### 2.5. Annual changes in zooplankton biomass in the Oyashio area

Oyashio waters provide the summer feeding ground for small pelagic fishes in the western North Pacific (Yatsu, 2019), and the zooplankton biomass in these feeding grounds is known to affect the population dynamics of these fishes (Odate, 1994; Tadokoro et al., 2005; Yatsu, 2019). Annual changes in mean zooplankton biomass in June and July in the Oyashio area from 1971 to the 1980s have been reported in previous studies (Odate, 1994; Tadokoro et al., 2005). In these studies, Oyashio waters were defined as the area where temperature at 100-m depth was less than 5 °C (Kawai, 1972), and mean monthly mesozooplankton biomass collected by plankton nets was calculated within this area. In June, the eastern margin of this area was bounded by the longitude 145°E from 35–37°N, and by the longitude 146°E from 38–43°N (Fig. 2). In July, the eastern margin was bounded by the longitude 155°E from 36–45°N. The area in July of water depth <500 m was excluded from this analysis. Although in these previous analyses mean mesozooplankton biomass was calculated without the use of a flow meter, a recent study of filtering-rate variability for vertical plankton tows has suggested that data lacking flow-meter information are still suitable for large-scale comparisons such as annual comparisons of zooplankton biomass (Takasuka et al., 2017). Therefore, we used these analysis results to compare mesozooplankton biomasses between the current and previous period of increase. For the current period of increase, mesozooplankton was collected in NORPAC and modified-NORPAC nets towed vertically from a depth of 150 m during the years 2011–2019. Both net types had the same mouth diameter (45 cm) and mesh size (0.335 mm), but were of different lengths and shapes. We did not correct for the difference in the nets because they have been reported to have the same sampling efficiency (Kotani, 1994).

We estimated mean monthly mesozooplankton biomass (g/m^2^) by calculating biomass per 150 m of depth within the defined areas to compare against the previous studies (Fig. 2). Biomass data for June were drawn from Odate (1994) and data for July were extracted from figures published in Tadokoro et al. (2005) by using WebPlotDigitizer v. 4.5 (Rohatgi, 2021).

### 2.6. Statistical analysis

Mean *r*_*1*_ by age was compared by using Tukey’s honestly significant difference (HSD) test. Relationships between *L*_*t*_ and *N*_*c*_ were analyzed by using general linear models to examine the density dependence of growth. We hypothesized that the strength of density dependence would differ between the previous and current periods of increase. Therefore, the effects of *period* (a categorical variable) and the interaction of *N*_*c*_ and *period* (*N*_*c*_ × *period*) were also examined in the model. We used *L*_*t*_ as the response variable, and *N*_*c*_, *period*, and *N*_*c*_ × *period* as explanatory variables. The significance of explanatory variables was examined by a likelihood-ratio test.

All statistical tests were performed in R v. 3.6.3 (R Core Team, 2020). VBGF parameters were estimated by using the package FSA (Ogle et al., 2020), and likelihood-ratio tests were conducted by using the packager car (Fox and Weisberg, 2019).

## 3. Results

A total of 1642 Japanese sardine of SL 82–245 mm were used for otolith analysis. The proportion of otoliths with an opaque zone on the outer margin of the posterior side was higher from May to September in all age classes (Fig. S1). Opaque and translucent zones were clearly visible, and ages were determined for all individuals (Fig. S2). The maximum estimated age was 7 years (SLs 225 and 230 mm) (Table 1). *r*_*1*_ values were measured for 348 samples. Mean *r*_*1*_ values for ages 0 to 4+ were 980.7 µm, 1079.7 µm, 1082.5 µm, 1088.8 µm, and 1106.5 µm, respectively, and the mean *r*_*1*_ for age 0 was significantly lower than that for the other ages (*p* < 0.001) (Fig. S3).

**Table 1.** Standard length (SL) and age (*t*) ranges, number of individuals (*n*), and estimated parameters (95% confidence interval [CI]) of von Bertalanffy growth formulas, asymptotic SL (*L*_inf_), growth coefficient (*K*), and hypothetical age at zero SL (*t*_0_), by year-class from the present study and modified from Wada & Kashiwai (1991a, 1991b).

| Source | Year class | $n$ | SL range (mm) | $t$ range (years) | $L_{inf}$ (95% CI) | $K$ (95% CI) | $t_0$ (95% CI) |
| --- | --- | --- | --- | --- | --- | --- | --- |
| This study | 2011 | 89 | 101–234 | 0.8–6.5 | 223.5 (217.0, 229.9) | 0.72 (0.60, 0.84) | −0.20 (−0.35, −0.04) |
|  | 2012 | 117 | 106–245 | 0.7–7.3 | 235.9 (230.1, 241.7) | 0.67 (0.60, 0.75) | −0.30 (−0.40, −0.20) |
|  | 2013 | 256 | 123–235 | 0.7–6.3 | 224.4 (217.7, 231.1) | 0.81 (0.65, 0.97) | −0.39 (−0.65, −0.14) |
|  | 2014 | 272 | 130–234 | 0.7–5.8 | 228.3 (220.8, 235.7) | 0.56 (0.47, 0.64) | −0.84 (−1.06, −0.63) |
|  | 2015 | 199 | 116–232 | 0.8–5.9 | 206.8 (202.3, 211.4) | 0.89 (0.72, 1.05) | −0.41 (−0.63, −0.19) |
|  | 2016 | 112 | 114–219 | 0.6–4.9 | 201.0 (193.5, 208.5) | 0.75 (0.54, 0.96) | −0.78 (−1.16, −0.40) |
|  | 2017 | 179 | 83–214 | 0.4–3.9 | 205.8 (185.5, 226.1) | 0.48 (0.29, 0.67) | −1.18 (−1.67, −0.69) |
|  | 2018 | 269 | 100–198 | 0.4–2.9 | 203.5 (185.4, 221.7) | 0.47 (0.32, 0.63) | −1.20 (−1.58, −0.81) |
| Wada & Kashiwai (1991a, 1991b) | 1976 | - | - | 1–4 | 225.0 | 0.53 | −1.11 |
|  | 1977 | - | - | 1–4 | 225.4 | 0.52 | −0.94 |
|  | 1978 | - | - | 1–4 | 212.4 | 0.65 | −0.88 |
|  | 1980 | - | - | 1–4 | 204.7 | 0.37 | −1.84 |
|  | 1981 | - | - | 1–4 | 211.5 | 0.32 | −2.50 |
|  | 1982 | - | - | 1–4 | 206.5 | 0.46 | −1.40 |

VBGF parameters were estimated for the 2011–2018 YCs from 1491 samples (Table 1). *L*_*inf*_ ranged from 201.0 (2016 YC) to 235.9 (2012 YC), *K* from 0.47 (2018 YC) to 0.89 (2015 YC), and *t*_*0*_ from −1.20 (2018 YC) to −0.20 (2011 YC) (Table 1). The estimated VBGFs of the most recent eight YCs showed lower growth in later cohorts (Fig. 3 and Table S1). Comparisons between the previous and current periods of increase indicated similar growth for the 1980–1982, 2017, and 2018 YCs (Fig. 3).

**Fig. 3.**
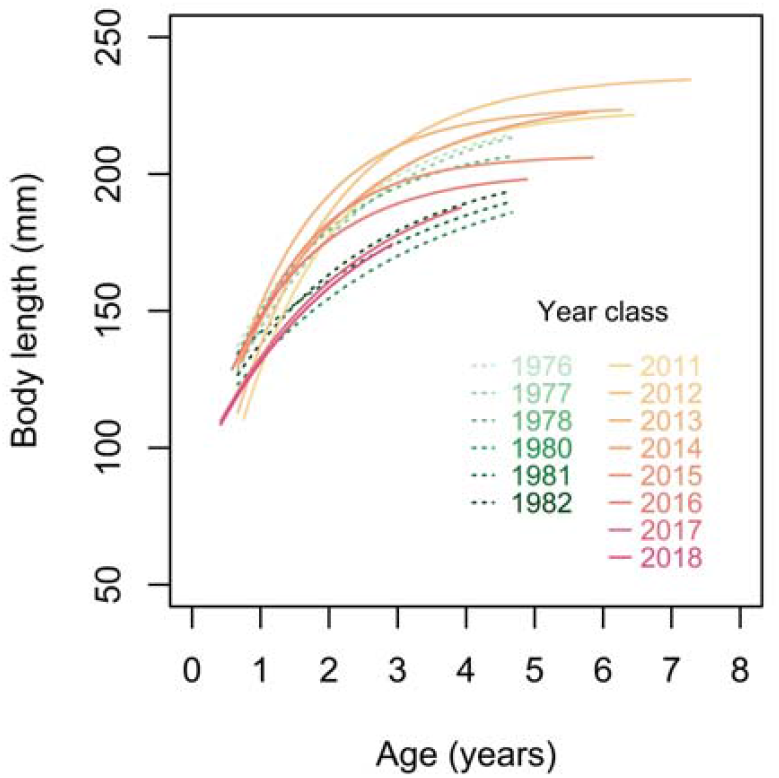
Comparison of von Bertalanffy growth formulas of Japanese sardine among 14 year-classes. Formulas from 1976 to 1978 and 1980 to 1982 are based on parameters modified from Wada and Kashiwai (1991a, 1991b).

The VBGFs estimated from otolith data showed roughly similar growth patterns to those estimated from scale data. However, the confidence intervals for the otolith data were wider (Table S1 and Fig. S4).

The general linear model and likelihood-ratio tests for growth and abundance in age 2 to age 4 sardine showed that *N*_*c*_ and *N*_*c*_ × *Period* were significant variables (Table 2). The predicted relationships between *L*_*t*_ and *N*_*c*_ had higher slopes during the current period of increase than during the previous period, indicating that the density dependence of growth has become stronger in recent years (Fig. 4 and Table 2).

**Table 2.** Results of a linear model with likelihood-ratio tests for growth of Japanese sardine. *N*_*c*_, cumulative sum of Japanese sardine abundance; *Period*, the previous or current period of increase.

| Age (*t*) | Likelihood-ratio test |  |  | Selected model |  |  |
| --- | --- | --- | --- | --- | --- | --- |
| | Variable | $\chi^2$ value | *P* value | Parameter | Estimate | SE |
| 2 | $N_c$ | 16.40 | <0.001 | Intercept | 191.413 | 3.173 |
| | *Period* | 0.05 | 0.820 | $N_c$ | −0.039 | 0.006 |
| | $N_c \times \textit{Period}$ | 10.61 | 0.001 | $N_c \times \textit{Period}$ | −0.121 | 0.028 |
| 3 | $N_c$ | 22.90 | <0.001 | Intercept | 214.617 | 3.503 |
| | *Period* | 0.22 | 0.640 | $N_c$ | −0.032 | 0.004 |
| | $N_c \times \textit{Period}$ | 10.98 | <0.001 | $N_c \times \textit{Period}$ | −0.092 | 0.022 |
| 4 | $N_c$ | 21.24 | <0.001 | Intercept | 227.255 | 4.198 |
| | *Period* | 0.05 | 0.817 | $N_c$ | −0.024 | 0.003 |
| | $N_c \times \textit{Period}$ | 5.97 | 0.015 | $N_c \times \textit{Period}$ | −0.068 | 0.012 |

**Fig. 4.**
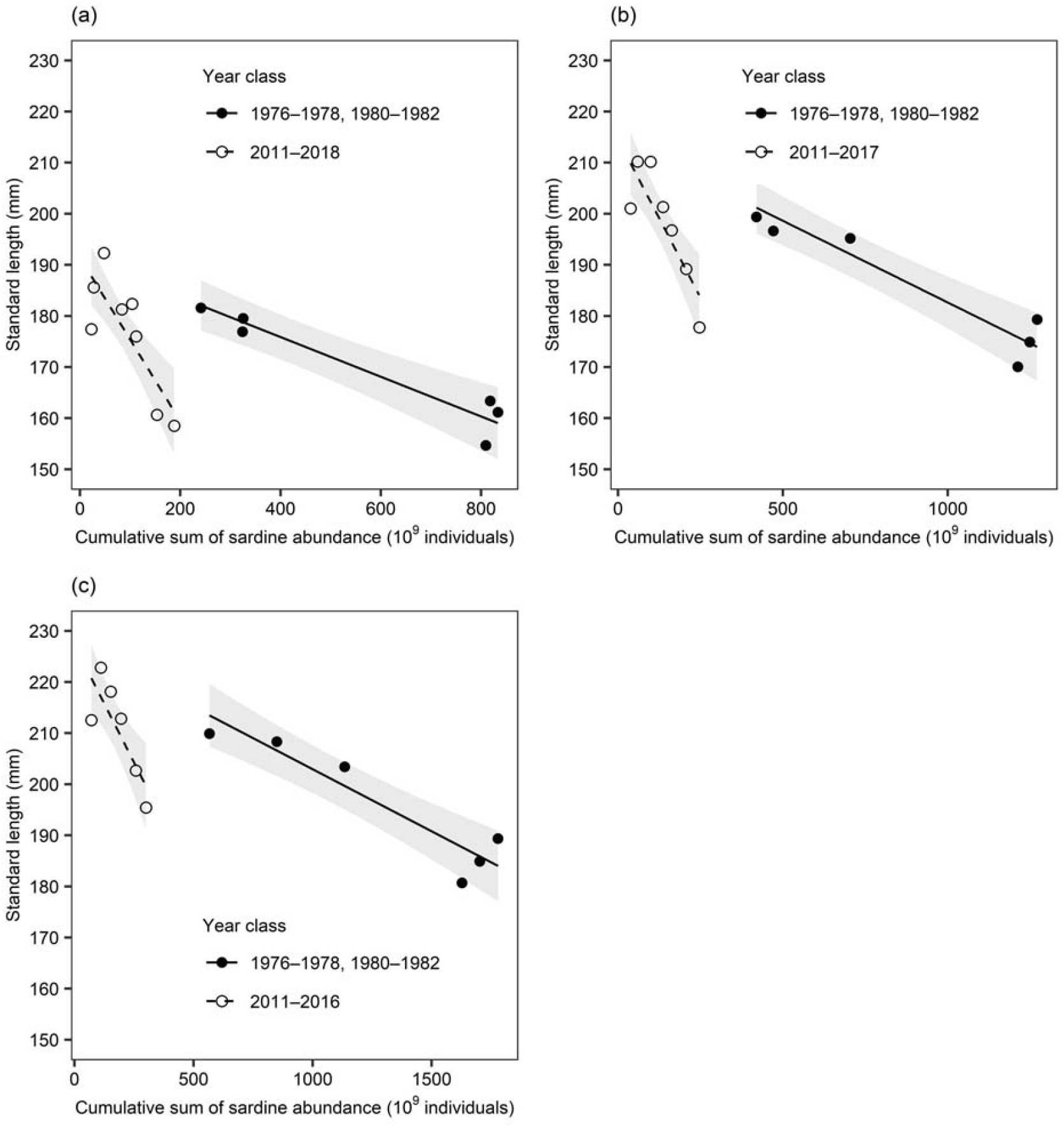
Relationship between the cumulative sum of Japanese sardine abundance and estimated standard length using von Bertalanffy growth formulas at ages (a) 2, (b) 3, and (c) 4. Circles and lines indicate observed values and predicted linear functions from the general linear model, respectively. Shaded areas show 95% confidence intervals.

Mean monthly mesozooplankton biomass from 1971–1985 ranged from 12.2 to 65.7 g/m^2^ in June and from 3.1 to 70.7 g/m^2^ in July (Fig. 5). Mean biomass declined from 1971 to the 1980s, likely due to a combination of climatic changes and the increase in Japanese sardine abundance after 1980 (Tadokoro et al., 2005). From 2011–2019, mean mesozooplankton biomass ranged from 27.3 to 57.8 g/m^2^ in June of each year and from 11.1 to 30.6 g/m^2^ in July of each year. In June, mean mesozooplankton biomass for 1971–1979 (43.0 g/m^2^) was similar to that for 2011–2019 (40.7 g/m^2^). In July, mean monthly mesozooplankton biomass for 1971–1979 (18.7 g/m^2^) was less than that for 2011–2019 (39.3 g/m^2^) and similar to that in the early 1980s, when Japanese sardine abundance was more than four-fold higher than currently.

**Fig. 5.**
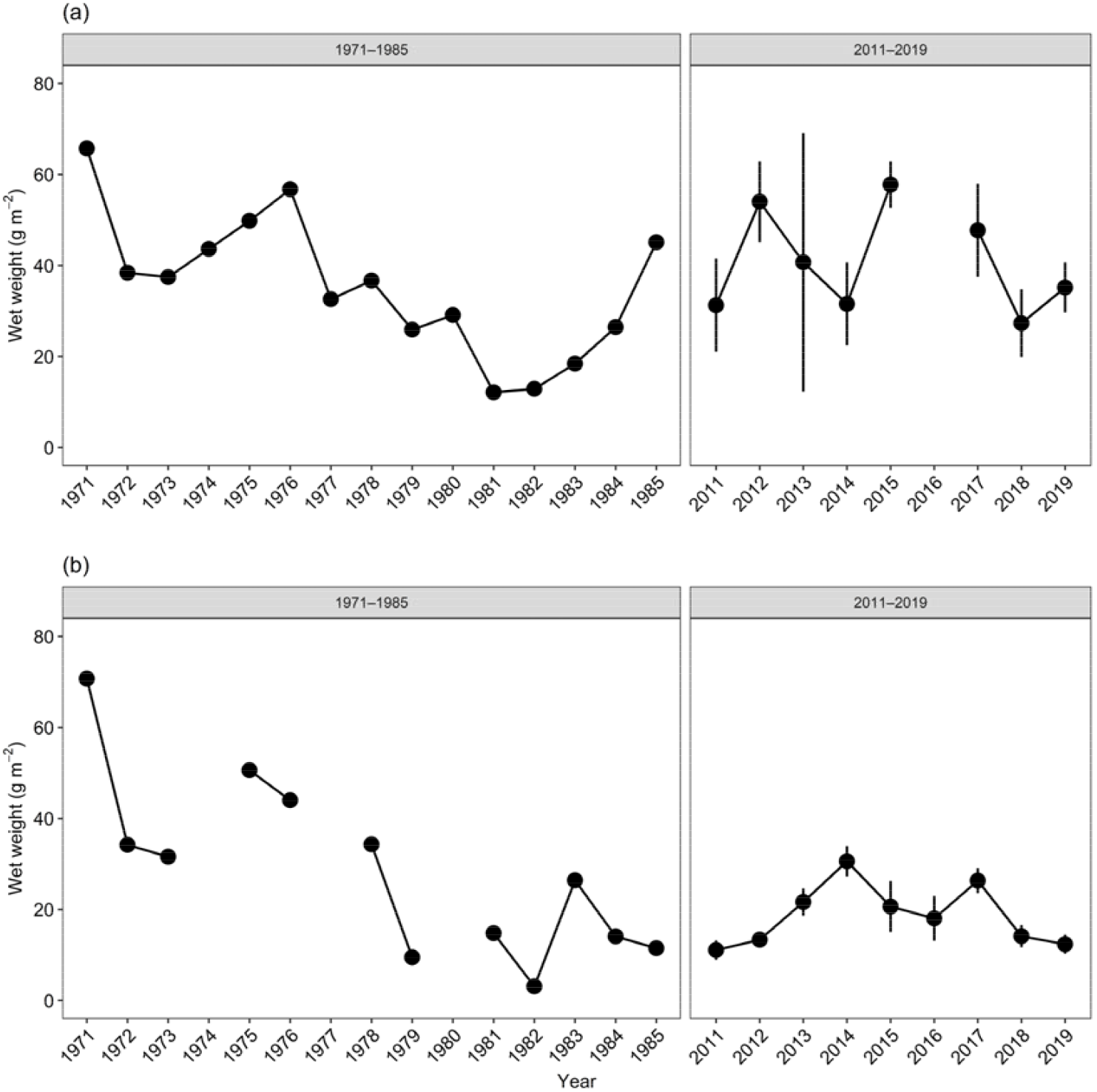
Annual changes in mean mesozooplankton biomass in the areas defined by Odate (1994) and Tadokoro et al. (2005) during (a) June and (b) July. Vertical bars indicate standard errors. June data from 1971 to 1985 are from Odate (1994), and July data are from Tadokoro et al. (2005).

## 4. Discussion

Our results show a negative relationship between length at ages 2 – 4 and the cumulative sum of abundance, which indicates that growth rates have a negative density dependence. Additionally, the strength of this density dependence appears to differ between the previous and current period of increase (Fig. 4). Although there have been several reports of negatively density-dependent growth among the Clupeidae, such as in the genera *Clupea* (Brunel and Dickey-Collas, 2010), *Sardinops* (Dorval et al., 2015; Kawabata et al., 2011; Ohshimo et al., 2009), and *Sardina* (Véron et al., 2020), these previous studies did not examine changes in the strength of density dependence over time. A study on Northeast Arctic cod (*Gadus morhua*) did suggest that the density dependence of growth changed across several decades in response to environmental factors such as water temperature, prey availability, and the status of the North Atlantic Oscillation, but the effects of these factors were not fully explored (Eikeset et al., 2016).

Food availability is one of the most important factors affecting the density dependence of growth (Amundsen et al., 2007; Jenkins et al., 1999; Kamimura et al., 2021). Our results show that mean mesozooplankton biomass in the Oyashio area in June was roughly equivalent in the 1970s (average, 43.0 g/m^2^) and 2010s (40.7 g/m^2^), but mean biomass in July was considerably lower in the 2010s (18.7 g/m^2^) than in the 1970s (39.3 g/m^2^) (Fig. 5). This suggests that lower food availability in summer feeding grounds during and after July has caused stronger density-dependent growth through increases in intra-specific competition during the current period of increase.

Japanese sardine feed on both zooplankton and phytoplankton (Yatsu, 2019), but feed more intensively on zooplankton in summer (Kawasaki, 1984). This is also consistent with the importance of mesozooplankton biomass during and after July in the Oyashio area for Japanese sardine growth. Changes in the spatial distribution of Japanese sardine could also affect the strength of density dependence by changing food availability and the energetic cost of feeding. Indeed, as Japanese sardine populations increase, the expansion of feeding grounds to offshore waters might affect the strength of density-dependent growth (Wada and Kashiwai, 1991b). Similarly, temporal changes in the density dependence of body condition in sprat (*Sprattus sprattus*) in the Baltic Proper have been suggested to have been triggered by spatial changes in habitat in response to predation pressure (Casini et al., 2014). Future studies of changes in the spatial distribution of Japanese sardine could provide further insight into this possible mechanism.

Both bottom-up and top-down effects are thought to impact annual changes in zooplankton biomass in the western North Pacific (Chiba et al., 2004, 2006; Odate, 1994; Tadokoro et al., 2005). In fact, reduced chlorophyll-*a* concentrations following the 1976/1977 climatic regime shift and the flourishing of Japanese sardine have been suggested as possible factors behind the decrease in summer mesozooplankton biomass in the 1980s (Tadokoro et al., 2005). During the current period of increase, although Japanese sardine abundance has been similar to or lower than that reported for the previous period of increase (Fig. 1) (Wada and Jacobson, 1998), mean mesozooplankton biomass in July has remained lower than during the previous period. This likely reflects the importance of bottom-up processes in limiting mesozooplankton biomass during recent years. SSTs from July to September in the western North Pacific have increased since 2010, and are currently 1–2 °C higher than in the 1980s (Kuroda et al., 2020). This temperature increase is a possible factor in the recent decline of mesozooplankton biomass.

Temporal changes in the abundance of chub mackerel (*Scomber japonicus*), an abundant and planktivorous small pelagic fish in the western North Pacific (Nakatsuka et al., 2010; Taga and Yamashita, 2018; Yatsu, 2019), provide further evidence of the importance of bottom-up processes during the current period. Mackerel abundances ranged between 18 × 10^9^ to 25 × 10^9^ individuals from 1970 to 1978, and then decreased to 8 × 10^9^ individuals in 1982 (Yukami et al., 2020). Since 2011, abundances have increased, and remained at similar levels during the years 2013 to 2017 (23 × 10^9^ to 28 × 10^9^ individuals) as those reached in the 1970s. In fact, abundances in 2018 (46 × 10^9^ individuals) and 2019 (35 × 10^9^ individuals) were the highest ever recorded. Although this might be expected to exert a negative top-down effect on mesozooplankton biomass, in reality there has been no clear relationship between mesozooplankton biomass and chub mackerel abundance since 2011 (Fig. 5). Follow-up studies on possible bottom-up effects and on intra- and inter-specific competition based on dietary analysis would help clarify changes in food availability for planktivorous fishes in this area.

Estimated growth trajectories for the 2017 and 2018 YCs were similar to those for the 1980 to 1982 YCs (i.e., the lowest-growth YCs since 1976) (Fig. 3), even though total Japanese sardine abundance and biomass since 2018 were only about one-third to one-fifth of that in the 1980s (Fig. 1). This suggests that the abundance of small pelagic fishes is already approaching the carrying capacity of the North Pacific. Observations of decreased chub-mackerel growth and body condition in the western North Pacific since 2013 also support this contention (Kamimura et al., 2021). Previous analyses have indicated that population increases of Japanese sardine have been stunted by low growth since the 1980 YC (Wada and Kashiwai, 1991b); therefore, sardine abundance might not increase as much in the near future as it did in the 1980s unless food availability improves.

Our results show that the strength of density dependence of growth differed across the study period. Although previous studies have examined the density dependence of Japanese sardine growth and egg production by using long-term multi-decadal datasets (e.g. Kawabata et al., 2011; Takasuka et al., 2019), such studies implicitly assumed that the strength of density dependence is time invariant. It is important when examining the relationship between biological characteristics and population abundance for the purpose of applying them to fisheries management and future forecasts to allow for temporal changes in these relationships, as has been indicated by previous reports of likely changes in the growth–recruitment relationship of this species over time (Furuichi et al., 2020b).

Our study is the first to assess the annual periodicity of otolith increments in Japanese sardine, and is also the first to successfully carry out otolith-based age determination and growth estimation in this species. Otolith increments were clearly distinguishable by the surface reading method (Fig. S2) and proved as useful for age determination in this species as has been the case for other sardines (Fletcher and Blight, 1996; ICES, 2019; Yaremko, 1996). The mean *r*_*1*_ of age 1+ individuals ranged from 1079 to 1108 µm, which is similar to mean *r*_*1*_ for *Sardina pilchardus* (about 1.1 mm; ICES, 2019). This result can be used to guide the identification of the first annual ring in Japanese sardine.

The mean *r*_*1*_ of age-1+ fish was about 100 µm larger than that of age-0 fish; in other words, the mean length of age-1+ fish when the first translucent zone was formed was about 10 mm longer than that of age-0 fish as calculated from the relationship between otolith radius and body length in juvenile Japanese sardine (Ohshimo et al., 1997). This suggests that larger Japanese sardine were more likely to survive their first year of life. This is consistent with previous reports that, in addition to larval survival, cumulative mortality between the egg stage and age 1 is important for recruitment success (Watanabe et al., 1995).

Otolith- and scale-based VBGFs showed similar growth trajectories, although the otolith-based VBGFs had wider confidence intervals simply due to the smaller sample size of length-at-age measurements (Fig. S4 and Table S1). This suggests that there is little difference among mean ages as determined by otolith and scale examination, and that the differences between the methods did not meaningfully affect our results. Further work is still needed to assess the precision and accuracy of age determination in this species from otolith- and scale-based methods.

## 5. Conclusions

Our study clarified changes in the recent growth of late-juvenile and adult Japanese sardine, and is the first to confirm the annual periodicity of otolith ring formation in this species. Otolith analysis is a powerful tool for understanding experienced environments and for reproducing migration histories of individual sardine (Sakamoto et al., 2017, 2019); therefore, our results should provide a strong basis for such research in the future. Our growth analyses showed that growth decreased with increasing abundance, and that density dependence in the current period of increase is likely to be stronger than in the past. Mean mesozooplankton biomass in the Oyashio area in July was also reduced in recent years as compared to the previous period of increase. Therefore, we suggest that the stronger density dependence in recent years is caused by lower food availability, and that zooplankton biomass during and after July in the Oyashio area is an important determinant of the strength of density dependence. The implied lower carrying capacity in the western North Pacific may indicate that Japanese sardine abundance will not increase to past levels. Similar considerations of environmental conditions could be useful for determining reference points for this stock, forecasting future stock status, and for constructing population dynamics models of this species.

## CRediT authorship contribution statement

Yasuhiro Kamimura: Conceptualization, Data curation, Methodology, Formal analysis, Investigation, Writing – Original Draft. Kazuaki Tadokoro: Data curation, Investigation, Resources, Writing – Review & Editing. Sho Furuichi: Data curation, Investigation, Resources, and Writing – Review & Editing. Ryuji Yukami: Data curation, Investigation, and Writing – Review & Editing.

## Acknowledgements

We thank R. Watanabe and Y. Niino for their help in otolith analysis. We appreciate the members of the Hachinohe Field Station at the Japan Fisheries Research and Education Agency and the crew of the T/V *Hokuho Maru* for their help in Japanese sardine sampling. This study was funded by the Japan Fisheries Research and Education Agency and the Fisheries Agency of Japan.

## Appendix A. Supplementary data

Supplementary material related to this article can be found in the online version.

