## Supplementary materials for "Stronger density-dependent growth of Japanese sardine with lower food availability: Comparison of growth and zooplankton biomass between a historical and current stock-increase period in the western North Pacific"

Table S1. Standard length (SL) and age (*t*) ranges, number of individuals (*n*), and estimated parameters (95% confidence interval [CI]) of von Bertalanffy growth formulas, asymptotic SL (*L*_inf_), growth coefficient (*K*), and hypothetical age at zero SL (*t*_0_), by year class estimated from scale-based age data.

| Year  class | *n* |  | BL range   (mm) | *t* range  (years) | *L_inf_* (95% CI) | *K* (95% CI) | *t_0_* (95% CI) |
| --- | --- | --- | --- | --- | --- | --- | --- |
| 2013 | 1496 |  | 93–253 | 1.0–7.2 | 218.3 (216.2, 220.3) | 1.08 (0.98, 1.18) | 0.07 (−0.02, 0.16) |
| 2014 | 4134 |  | 43–243 | 0.0–6.8 | 209.0 (207.6, 210.4) | 0.82 (0.79, 0.86) | −0.49 (−0.53, −0.45) |
| 2015 | 2717 |  | 58–238 | 0.2–5.9 | 215.9 (213.4, 218.5) | 0.61 (0.57, 0.65) | −0.94 (−1.03, −0.85) |
| 2016 | 2473 |  | 68–233 | 0.3–4.9 | 213.4 (210.6, 216.2) | 0.64 (0.60, 0.69) | −0.80 (−0.88, −0.72) |
| 2017 | 2604 |  | 43–238 | 0.3–3.9 | 209.8 (203.9, 215.7) | 0.50 (0.44, 0.57) | −1.20 (−1.37, −1.04) |
| 2018 | 2656 |  | 53–208 | 0.2–2.9 | 178.8 (176.0, 181.6) | 0.92 (0.83, 1.01) | −0.61 (−0.70, −0.52) |

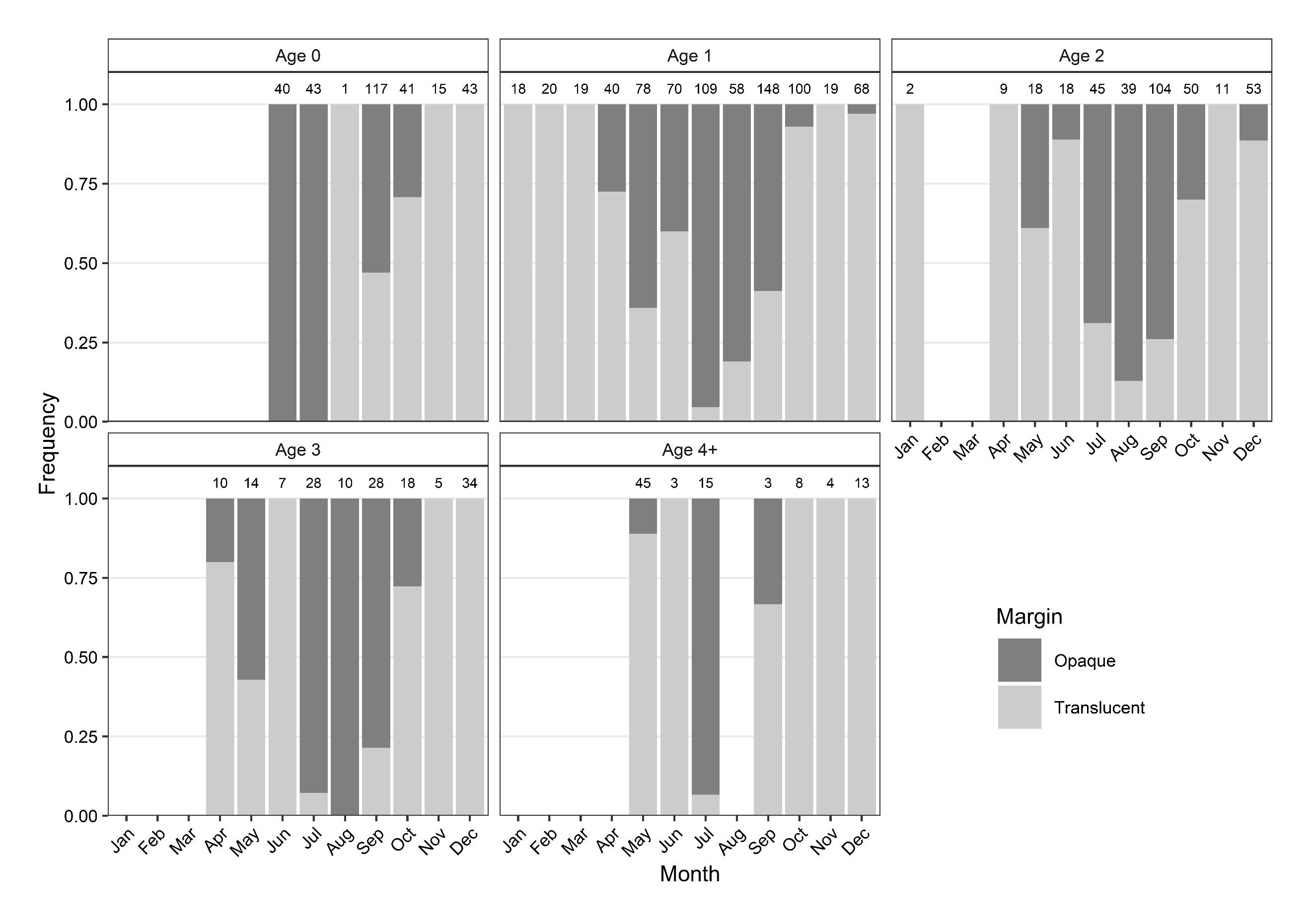

Fig. S1. Monthly changes in occurrences of opaque (dark grey) and translucent (light grey) zones at the outer otolith margin for fish of ages 0 to 4+. Numbers above bars denote sample sizes.

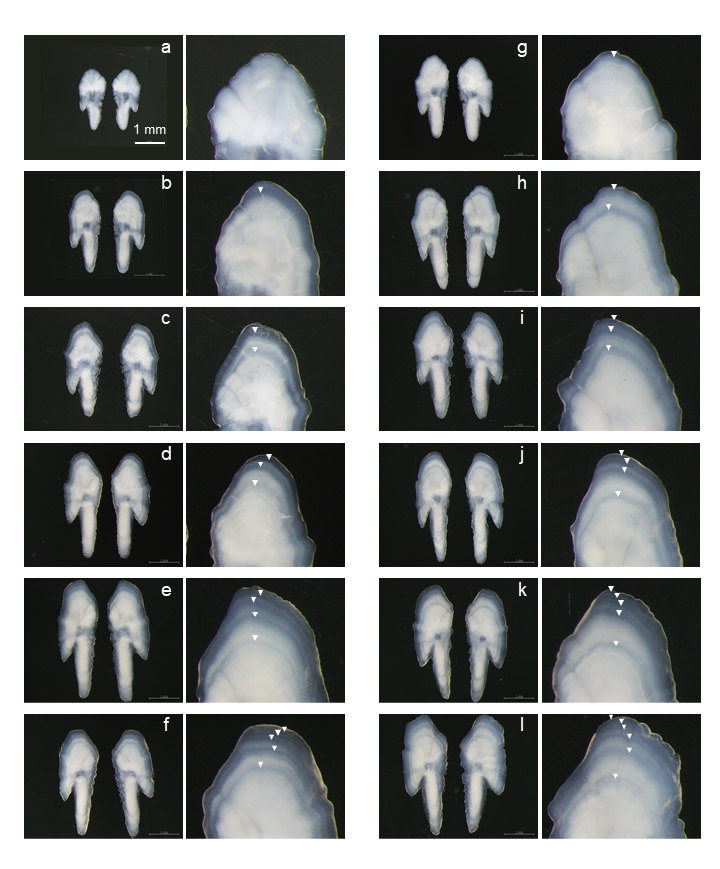

Fig. S2. Japanese sardine otoliths collected in July (a–f) and December (g–l). Left-hand panels show the entire otolith, and right-hand panels show enlarged views. Triangles indicate translucent zones. Years of collection, standard lengths, and determined ages are as follows: (a) 2019, 102.5 mm, age 0; (b) 2019, 147.3 mm, age 1; (c) 2019, 171.5 mm, age 2; (d) 2019, 193.1 mm, age 3; (e) 2019, 198.5 mm, age 4; (f) 2020, 197.0 mm, age 5; (g) 2015, 137.3 mm, age 0; (h) 2015, 172.0 mm, age 1; (i) 2016, 198.0 mm, age 2; (j) 2016, 220.0 mm, age 3; (k) 2017, 225.0 mm, age 4; and (l) 2016, 234.0 mm, age 5.

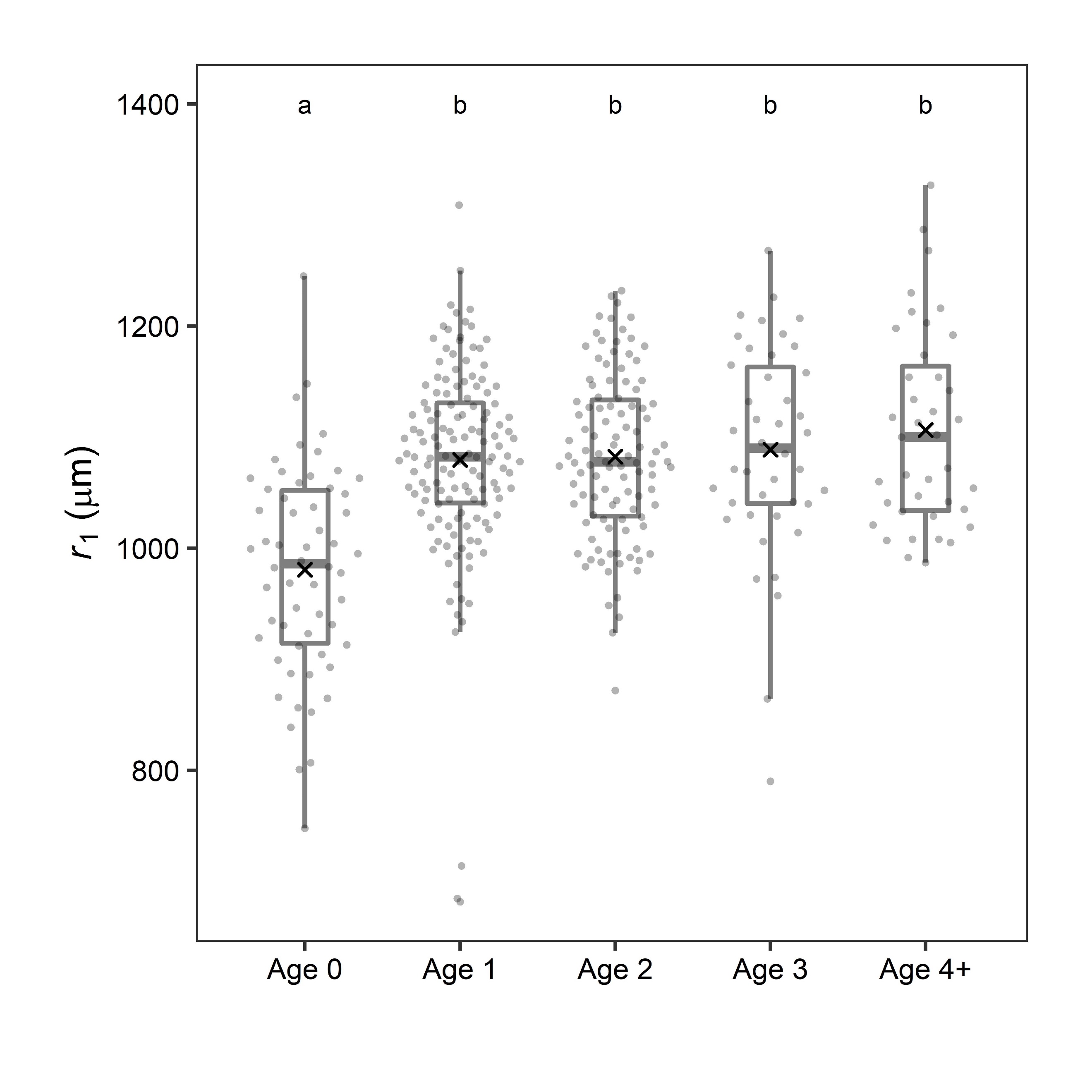

Fig. S3. Box plot of the otolith radius from the nucleus to the first translucent zone on the posterior side (*r_1_*). Line within boxes, upper and lower ends of boxes, and whiskers show the median, first and third quartiles, and ±1.5 times the interquartile range, respectively. Grey dots show raw data, and crosses indicate mean values. Age groups with different letters above the bars are significantly different at *P* < 0.001 (Tukey’s HSD).

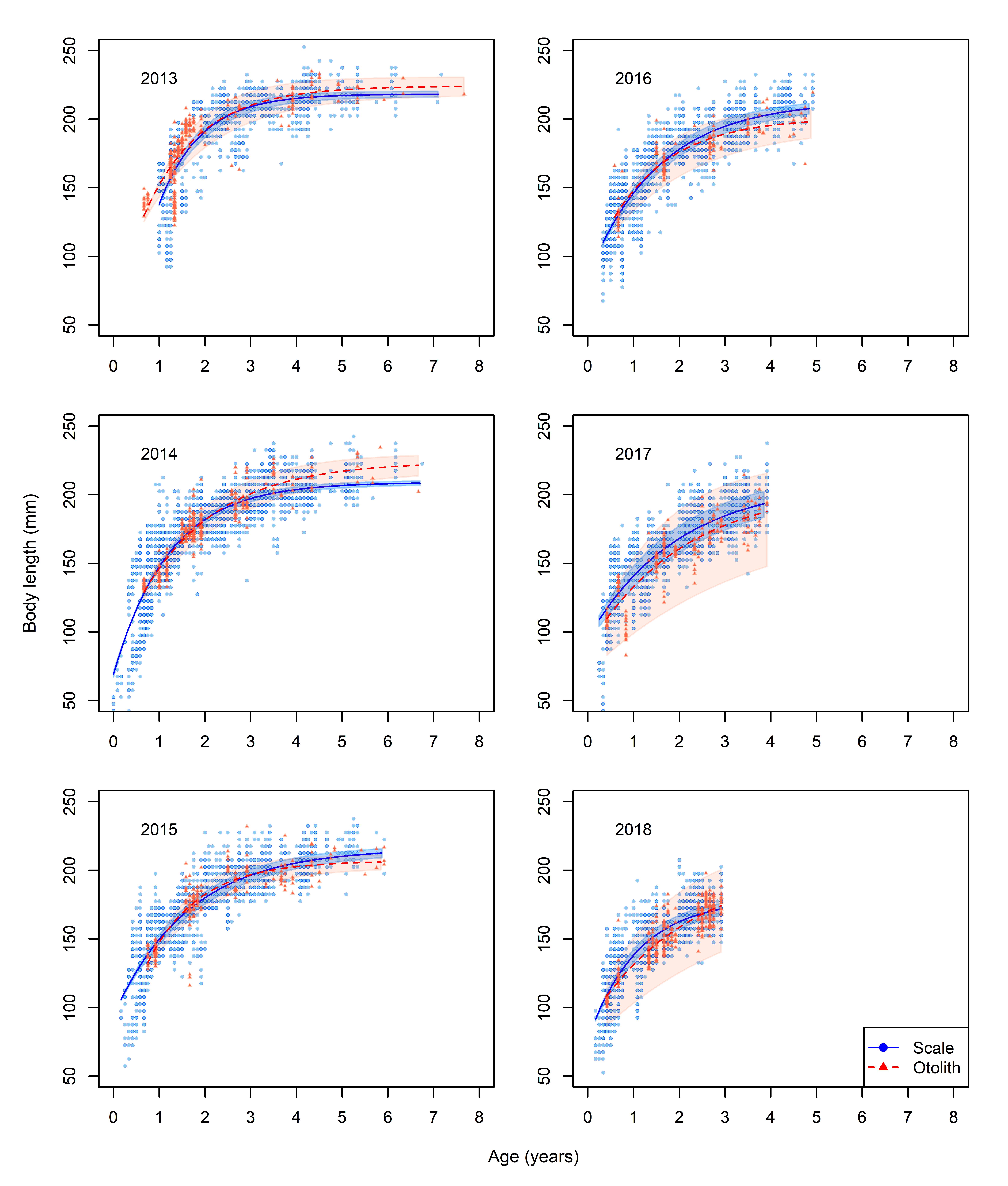

Fig. S4. Comparison of von Bertalanffy growth trajectories estimated by using two different aging characters for six year-classes. Red dashed lines and triangles show trajectories estimated from otolith-based aging data, and blue solid lines and circles show those estimated from scale-based aging data. Shaded areas represent 95% confidence intervals.
